# Small proteins from prokaryotes in marine water column at full ocean depth

**DOI:** 10.1101/2024.11.08.622638

**Authors:** Qing-Mei Li, Li-Sheng He, Yong Wang

**Author notes:** **Corresponding authors:** Yong Wang, PhD, Institute for Ocean Engineering, Shenzhen International Graduate School, Tsinghua University, Shenzhen, 518055, P. R. China, **E-mail:**, Li-Sheng He, PhD, Institute of Deep Sea Science and Engineering, Chinese Academy of Sciences, Sanya, Hainan, 572000, P. R. China, **E-mail:**.

## Abstract

Small proteins (SPs, *≤* 50 amino acids) are usually ignored by automated bioinformatic algorithms and difficult to be detected by traditional protein purification. In this study, 193,281 prokaryotic SP clusters were generated using 433,311 small open reading frames (sORFs) predicted in assemblies of 71 full-ocean-depth marine water metagenomes from the western pacific. Further filtration with at least 5 copies per million reads resulted in 75,581 prevalent SP clusters, among which 4,307 clusters have coding capacity as predicted by RNAcode. For these RNAcode-filtered SP (RfSP) clusters hosted largely by Proteobacteria, Thaumarchaeota and Marinimicrobia, 29.16% of them show homology () in all prokaryotic genomes. About 2% of the RfSP clusters were estimated as potential antimicrobial peptides and were distributed across all depth zones. Transcription of 7.96% of the RfSP clusters was verified in 12 deep-sea metatranscriptomes, and translation of 34 prevalent SP clusters was detected in 14 metaproteomes. The transcribed RfSP clusters are involved in processes such as DNA stability under stress condition, betaine transport and cell division regulation for a rapid response to varying deep-sea environments. This study uncovers a vast reservoir of SPs with pivotal functions in potential antibiotic activities, hyperoxide detoxification and pressure resistance for fitness of marine prokaryotes in deep ocean.

## Introduction

Small proteins (SPs) generally defined to be less than 50 amino acids (aa) have been detected in most organisms ^1,2^. Detecting SPs is still technically challenging ^3^, and thus more were identified by bioinformatic algorithms combined with multi-omics and ribosome sequencing with mass spectrometry ^4,5^. In bacteria, SPs participate in cellular morphogenesis, regulation of membrane-bound enzymes, and regulatory networks of protein kinase and signal transduction ^6,7^. SP also functions as anti-microbial proteins (AMPs), such as the lantibiotics contain 19–38 amino acids (<5 kD)^8^. These previous studies indicate that SPs are important in microbes due to more specific associated functions when adapt to environmental stresses, such as YobF in response to heat shock and MgrB, a 47-amino-acid membrane protein that is part of the PhoQ/PhoP regulon ^9,10^. A recent study of SPs in human microbiomes identified >4,000 SPs with transcription and translation evidence, most of which are novel and conserved ^11^. In addition, thousands of small and novel genes were predicted in global phage genomes ^12^.

The global oceans are the largest habitat for prokaryotes on Earth with an estimate of 5.5×10^29^ cells in the various marine environments ^13^. More than 99% of prokaryotes in oceans were uncultivated ^14,15^. Lacking cultivated representatives of deep-sea dominant prokaryotic species poses an obstacle to studying proteins with unknown functions, including novel bioactive SPs. More than 30,000 natural products such as peptides have been obtained from the deep ocean ^16,17^. In deep ocean (the part of the ocean below 200 meters), organisms may secret SPs with antitumor, antihypertensive and anticoagulant effects ^18,19^. Deep-sea zones host a huge number of viruses to predate microorganisms for rapid circulation of nutrients among the ecosystem ^20–22^. There are perhaps novel antiviral and bio-toxic SPs produced by deep-sea microbes for survival; however, due to this extreme sampling challenges, antiviral agents derived from deep-sea microbes are limited ^23^. A comprehensive survey of conserved deep-sea SPs can discover novel candidate bioactive SPs, and elucidate the mechanisms for adaptation of deep-sea microbes to extreme environment.

In recent years, genomes and ecological functions of microbiomes in the deepest water layer of hadal zone (>6,000 m) have been studied ^24–26^, as well as in the upper layers (0-6,000 m) ^27–29^.The increasing number of metagenomic datasets from the water column provides an unprecedented opportunity to explore SPs from marine water microbiomes. In this study, we identified 75,581 prevalent SP clusters from assemblies of 71 metagenomes of full-ocean-depth water samples collected from the South China Sea (SCS) and Mariana Trench (MT). Our analysis showed that 4,307 of these SP clusters were potentially transcribed and conserved across different phyla. Transcription and translation of certain conserved SPs were confirmed by searching against deep-sea metatranscriptomes and metaproteomes. This is the first step towards understanding the functions of SPs as a strategy used by microbes living in deep oceans, and it also remarkably broadens the potential sources of bioactive substances.

## Results and discussion

### Identification of prokaryotic small proteins from full-ocean-depth marine water samples

We obtained 69 metagenomes with 684 Gbp Illumina sequencing reads for the marine water samples that were collected from the South China Sea (n=40) and Mariana Trench (n=29) with the sampling depth ranging from 100 m to 10,911 m (Fig. 1 and Table S1). Two extra metagenomes were recruited from NCBI for the Mariana water column at depths of 0 m and 203 m (Table S1). The metagenomes were assembled into contigs individually, which resulted in 42,792,862 contigs. A total of 433,311 short open reading frames (sORFs) with *≤*50 amino acids were identified from 42,706,554 prokaryotic contigs (Fig. S1). To reduce computational cost, these sORFs were further clustered into a dataset consisting of 193,281 SP clusters with pairwise identity and length coverage thresholds of 50% and 95%, respectively. To remove rare SPs, we retained the SP clusters with *≥* 5 copies per million metagenomic reads (CPM) for at least one metagenome, and this resulted in 75,581 prevalent SP clusters. Among 75,581 SP clusters, containing *≥*3 SP members and were not the last ORF of the respective contigs, 4,307 of them were able to encode an SP as predicted by RNAcode ^30^, which accounted for ∼5.70% of the prevalent SP clusters. The representative sequence of these RNAcode-filtered SP was named as RfSP in the following content description. 87.09% of representative sequences of RfSP clusters do not have a homolog in 80 non-marine metagenomes, including human associated microbiomes, bioreactors and lakes (Table S2). Similarly, 78% of the human RfSP clusters could not be mapped to non-human metagenomes ^11^. The difference may be due to larger database (5,829 samples) used for the analysis of human microbiome-encoded SPs. An alternative interpretation is that most of RfSPs encoded by the prokaryotes collected from the remote Mariana Trench were rarely encoded by the non-marine microbiomes.

**Figure 1.**
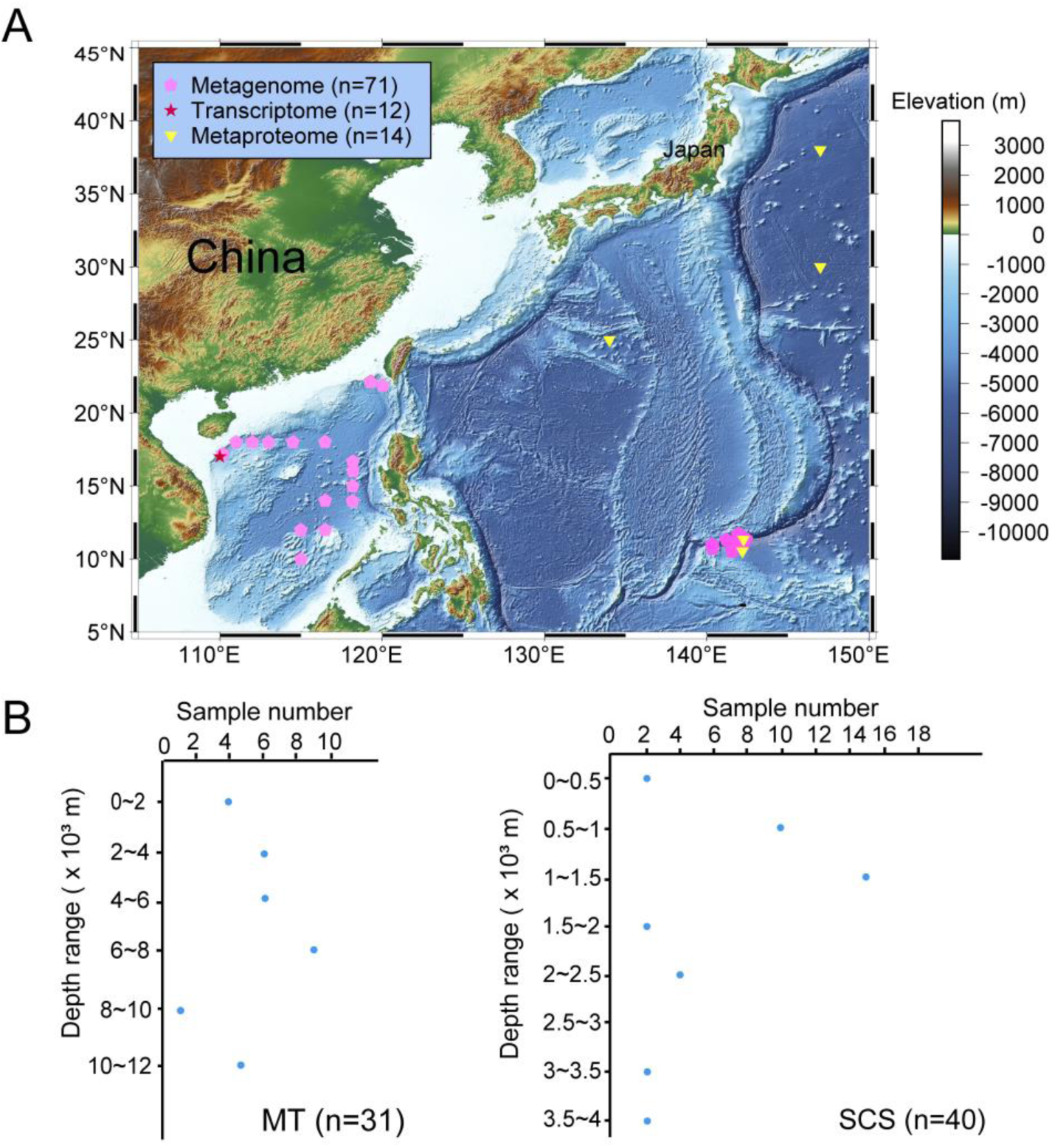
Sampling sites and vertical distribution of the marine water samples collected from the South China Sea (SCS) and the Mariana Trench (MT). (A) Pink pentagons represent the water samples obtained from the sites in the South China Sea and the Mariana Trench for metagenomes. The sampling sites for metatranscriptomes and metaproteomes are indicated by a red star and yellow triangles, respectively; (B) Number of samples collected from different depth ranges.

### Characteristics of small proteins from the full-ocean-depth samples

The 4,307 RfSP clusters in this study were largely longer than 40 aa (70.49% falling in 45∼50 aa size range) (Figs 2A and S2), which is similar to those from the human metagenomes ^11^ and complete prokaryotic genomes ^31^. One of the key factors affecting transcriptional efficiency of the sORFs encoding the RfSP clusters is the ribosomal binding sequence (RBS) motif, which contains Shine-Dalgarno (SD) sequence and downstream 5∼10 bp spacers (Fig. S3A) ending at the start codon ^32^. sORF members without an RBS motif occurred in 19.04% of the RfSP clusters (Fig. S3B), corresponding to ∼23% of the genes in bacterial genomes without an RBS motif ^32^. Perhaps due to low sequencing coverage for some of our sORFs, their RBS motif was missing or truncated. Nevertheless, it has been reported that some genes without an RBS are able to be transcribed ^32–34^. Our evaluation of conservation degree of the RBS motifs showed that 8.56% of all the RfSP clusters have the same RBS motifs in ≥ 90% of the corresponding sORF members (Fig. 2B). This indicates that the RBS motifs in the RfSP clusters were highly diversified. We compared SD sequences (n=55) in the RBS motifs for the 4,307 RfSP clusters and those (n=55) of SP clusters from the human microbiomes ^11^, which identified 51 shared types (Table S3). Four types of SD sequences in the RBS were specific to the marine RfSPs. The five most frequent SD sequences in upstream of 66,934 sORFs of the RfSP clusters were 5’GGA/GAG/AGG’3, 5’GGAG/GAGG’3, 5’AGGAG’3, 5’TAA’3, and 5’AGGA’3 (Fig. 2C). Except for 5’TAA’3, the AG rich SD sequences have been previously reported in prokaryotic genomes ^32^. In addition, the AT-rich RBS motifs have been shown in *Bacteroides* ^35^.

**Figure 2.**
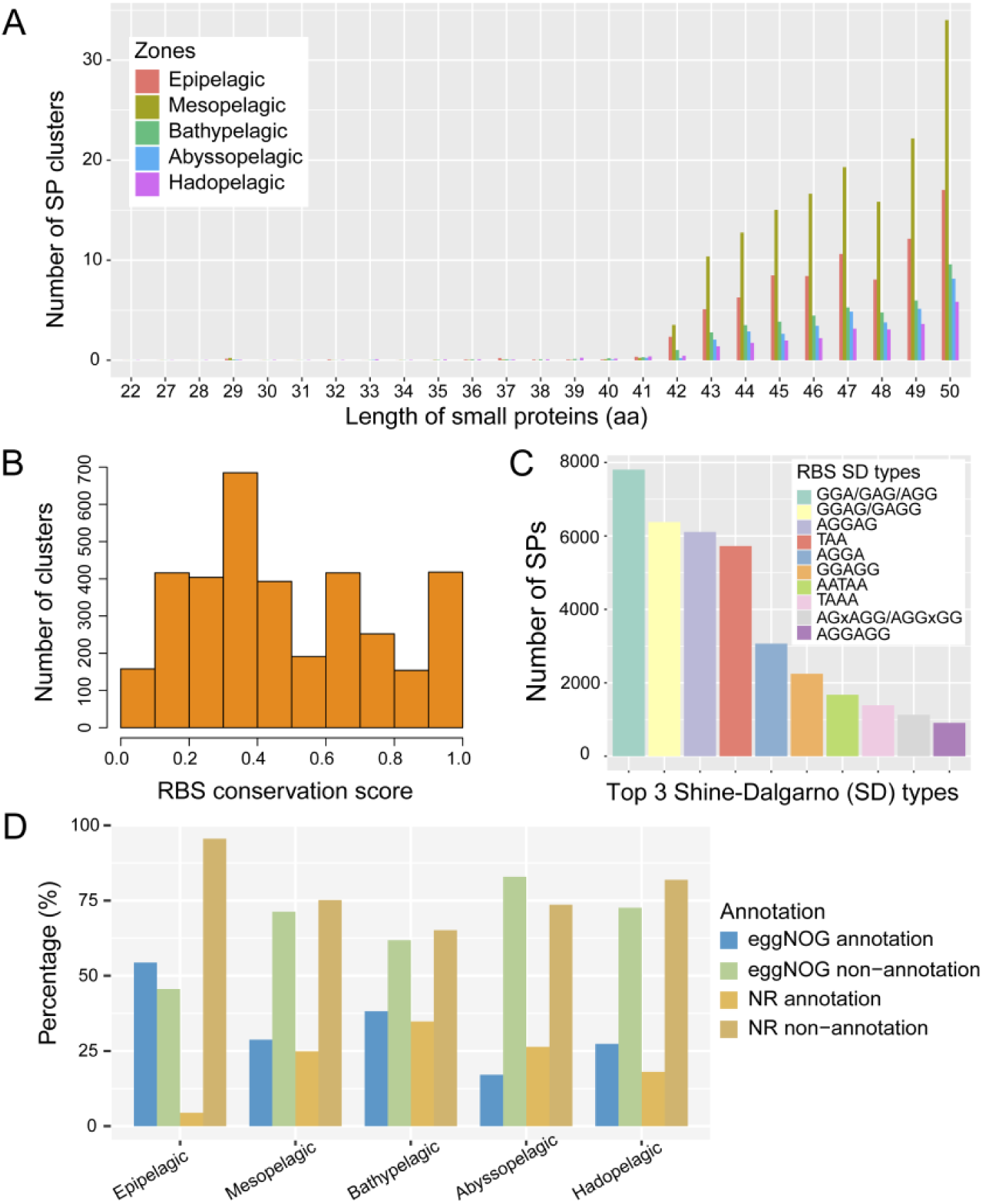
Characteristics of RfSP clusters from full-ocean-depth prokaryotes. (A) Size ranges of RNAcode-filtered SP clusters from different marine water column zones; the number of SP clusters in the depth zones was normalized by sequencing depth (Gbp) of metagenomes from the respective zones. (B) Distribution of RfSP clusters within different conservative score ranges of ribosomal binding site (RBS) (number of SP members characterized as frequently used RBS types divided by the total of SP members of a cluster) (Sberro, H., *et al*.); (C) Number of SP clusters with RBS motifs in the 4,307 RfSP clusters and the top 10 most frequently used Shine-Dalgarno (SD) sequences in these motifs; (D) Annotation results of SP clusters against EggNOG and NCBI_NR databases.

### Prediction of SP function

About 36% (n=1,534) of the 4,307 RfSP clusters have a homolog with *≥*90% length coverage and *≥*60% similarity from EggNOG and NCBI_nr databases (Fig. 2D). To examine the difference between the RfSPs and their homologs, we used the representative sequences of the RfSP clusters to search homologs in all available prokaryotic genomes and proteins of NCBI RefSeq. The search identified a homolog for 29.16% of the RfSPs. A total of 17.36% (213/1,227) RfSP clusters associated with an RBS motif were shorter than their homologs, most of which are much longer than 50 aa (Fig. S4). This indicates that some of the RfSPs are likely a truncated version of the homologous proteins essential in the activities of microorganisms for deep-sea adaptation. Previous studies showed that short versions of membrane proteins ^36^ and viral protein ^37^ are functional as well. When representative sequences of the RfSP clusters were blasted against to reviwed protein from UniPort database with less than 200aa, only 4.48% (193/4,307) of them has a hit with similar length (±10%). There were seven RfSP clusters were annotated as putative peroxiredoxin bcp, three were ferredoxin-2, three were cytochrome c oxidase subunit 4, two were cold shock protein A, two were ferric uptake regulation protein and so on (Table S4).

We expanded the annotation to all the 75,581 prevalent SP clusters. Conserved domains in these SP sequences were queried to CDD (conserved domain database)^38^, which showed that 17.79% (13,445/75,581) of the representative sequences could be assigned to known domains. The SPs were assigned with conserve domain more frequently descripted as phosphatidylinositol phosphate kinase (n=27), antitoxin component (n=46), IS3 family transposase (n=43), component of ABC-type branched-chain amino acid transport system (n=35), Bcp (Peroxiredoxin) (n=33), transposase protein (n=32, except IS3 family transposase) and so on (Table S4). Our result indicates that more than 80% of the SPs from marine waters were currently unknown (without conserved domain assigned) and need future efforts to examine their functions.

### Taxonomy and conservation of the small proteins

Using sequence-similarity-based classification software CAT ^39^, we revealed 38 phyla and 437 species in the taxonomic classification of the 4,307 RfSP clusters (Table S5). 20.36% and 52.52% of SP clusters could not be sorted to prokaryotic phylum and species levels, respectively (Fig. 3A). Proteobacteria and Thaumarchaeota are the most frequent bacterial and archaeal phyla, respectively, assigned to the SPs (Fig. 3B and Table S5); the SPs were also abundantly assigned to Marinimicrobia, Gammaproteobacteria and Gemmatimonadetes (Fig. 3C and Table S5), followed by three SAR11 species that were known to be dominant in marine waters ^40^. These most frequent phyla and species assigned to SP-bearing contigs have been previously reported in full-ocean-depth marine water samples ^41,42^, but functional SPs have not been reported yet.

**Figure 3.**
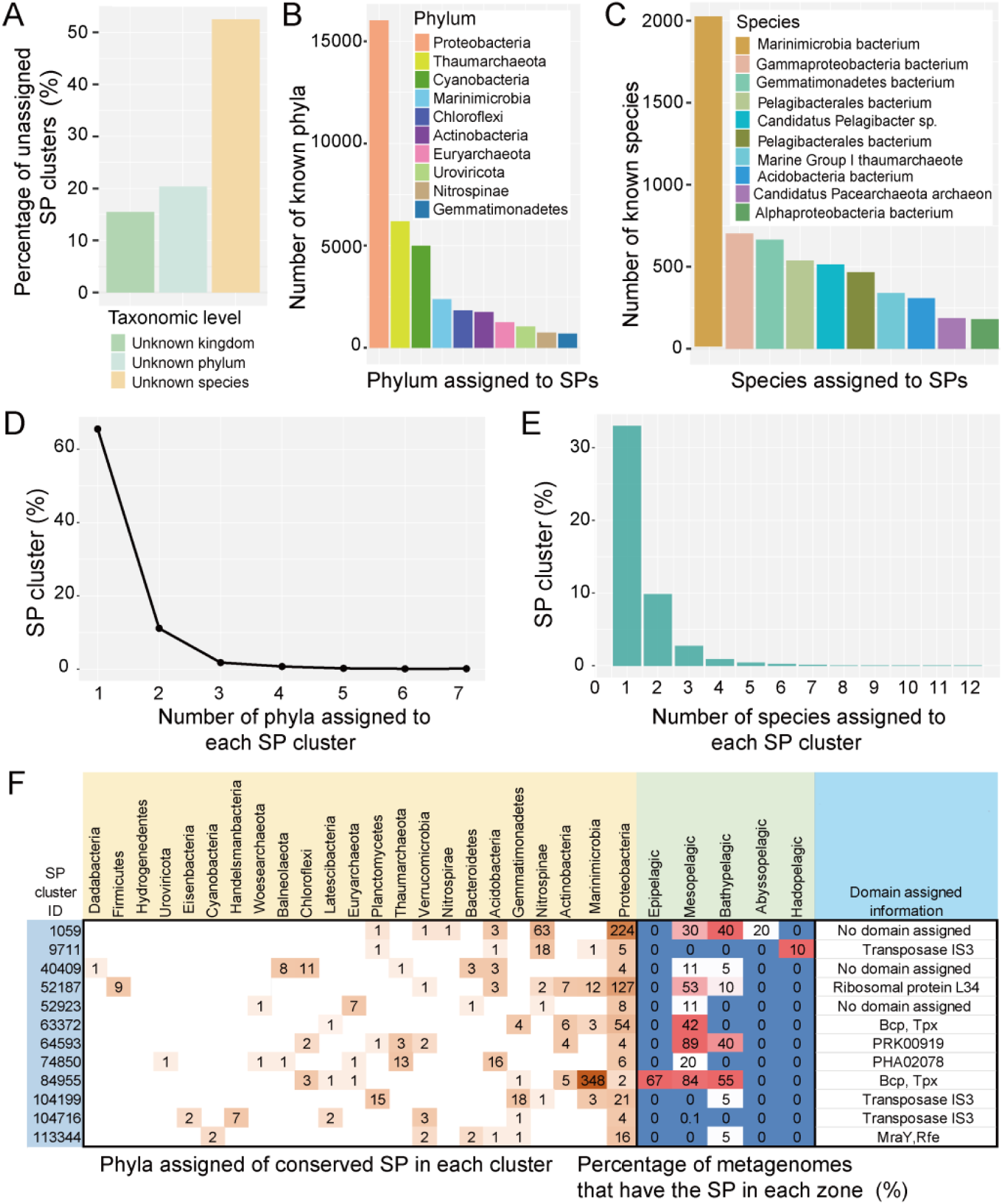
Taxonomy and conservation of RfSP clusters in marine prokaryotes. (A) The percentage of RfSP clusters without a taxonomic assignment at the kingdom, phylum and species levels according to CAT classification of the RfSP-encoding contigs; the top 10 most frequent phyla (B) and species (C) taxonomically assigned to RfSPs. Distribution ranges of RfSP clusters in different phyla (D) and species (E); (F) Taxonomy, depth zone distribution and domain function of 15 most widely spreading SP clusters. The RfSP members assigned to the phyla were counted and shown.

Distribution of SP clusters at phylum and species levels showed that 65.57% (2,824/4,307) of the RfSP clusters were encoded by one phylum (Fig. 3D) and 33.99% (1,421/4,307) of them were restricted in one species (Fig. 3E). To learn functions of widely distributed SP clusters, we examined RfSP clusters present in at least five phyla (Table S5 and Fig. 3F), which resulted in 15 SP clusters and most present in mesopelagic and bathypelagic zones (Fig. 3F). The RfSP cluster 84955 encoded by seven phyla is composed of RfSPs homologous to bacterioferritin comigratory protein (BCP), which is probably a new member of alkyl hydroperoxide peroxidase C family that acts on linoleic acid ^43^. This SP cluster was almost all derived from Marinimicrobia inhabiting epipelagic, mesopelagic and bathypelagic zones (Fig. 3F) and, remarkably, ∼84% of the mesopelagic metagenomes harbor the BCP-like RfSPs. The BCP (156 aa) of *Escherichia coli* is a versatile peroxiredoxin that catalyzes reduction of hydroperoxides and multiple substrates to confer resistance to oxidative stress ^44^. The RfSP cluster 63372 is the second BCP group. These RfSP BCPs might be assigned as Tpx (thiol peroxidase) as well, which is likely to reduce the peroxidase in eukaryotic Prx systems ^45^. Multiple sequence alignment of the BCP RfSPs and *E. coli* homologs showed that several amino acid positions were conserved with sequence identity being 51.16% and 44.44%, respectively (Fig. S5). The three-dimensional structure of the proteins was predicted using Swiss-model online, a total of 667 and 561 templates were found to match the RfSP clusters 84955 and 63372, respectively. The proteins most similar to RfSP clusters 84955 and 63372 were A0A413WBK2.1 and A0A382ARR4.1, respectively, based on the AlphoFold prediction (Fig. S6). The confidence level of the prediction was 0.97 and 0.92 for the RfSP clusters 84955 and 63372, respectively, in reference to the GMQE (Global Model Quality Estimate) (the GMQE values close to 1 mean higher reliability) ^46^. The A0A413WBK2.1 was thioredoxin-dependent thiol peroxidase and the A0A382ARR4.1 was alkyl hydroperoxide reductase subunit C/thiol specific antioxidant domain-containing protein. This indicates that RfSP clusters 84955 and 63372 are comprised of antioxidants. We showed that the SP cluster 63372 was restricted in mesopelagic zone and ∼79.41% of the members were probably affiliated with Proteobacteria (Fig. 3F). The employment of BCP SPs in these deep-sea prokaryotes for hyperoxide detoxification was probably stemmed from potentially intensive damages of superoxide on enzymes, DNA and lipids for deep-sea prokaryotes ^47,48^. Three widely distributed SP clusters 9711, 104199 and 104716 from the metagenomes were annotated to be transposase IS3 (Fig. 3F) that can mediate horizontal gene transfer (HGT) as the third class of mobile genetic elements ^49^.

### Identification of small proteins that may carry out antiviral and antimicrobial defense

About 11.26% (485/4,307) of the RfSP clusters or their up-/down-stream 10 proteins contain at least one member homologous to known defense proteins descripted in previous study ^50,51^. We identified four SP clusters with members being embedded and overlapping Clustered Regularly Interspaced Short Palindromic Repeats (CRISPR) associated genes such as an ancillary gene of type IIIB CRISPR-Cas system (*corA*) (Table S6) Among 165 members of the RfSP cluster 122274, 34.54% (57/165) are within a CRISPR-Cas system (Table S6). Particularly, nine conserved SP members overlap with different CRISPR-associated genes: TIGR00383 (*corA*), TIGR01587 (*cas3*), cd09639 (*cas3*_I), cd17930 (coding for DEXH/Q-box helicase domain of Cas3) and COG1199 (*dinG*) (Fig. 4 and Table S6). Cas3 is important in repelling invading genetic elements by serving as translocase and nuclease ^52,53^. In addition, 119 of the 165 members in RfSP cluster 122274 were classified into Thaumarchaeota (Table S6). The overlap of the RfSPs and the CRISPR associated genes can indicate an unknown strategy employed by Thaumarchaeota and other species to protect against viruses in marine water column.

**Figure 4.**
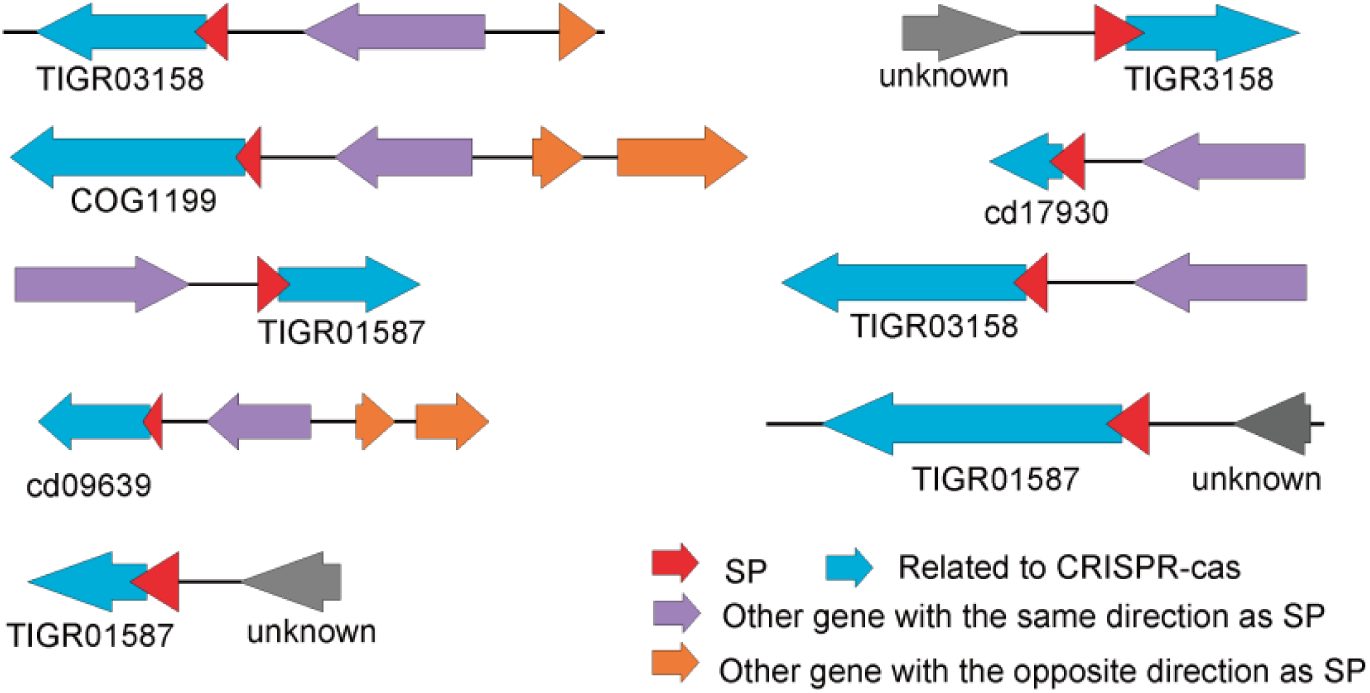
SP cluster involving CRISPR-Cas system. Nine SP sequences in SP cluster 122274, which are assigned to Thaumarchaeota, are adjacent to an ORF associated with a CRISPR-Cas system.

To *in silico* identify AMPs in the full-ocean-depth marine water column, the annotation of RfSPs against DBAASP and CAMPR3 databases showed that 2.11% (91/4,307) of the RfSP representative sequences were potential AMPs. Taxonomic classification of contigs that encode all the SP sequences of 91 potential AMP RfSP clusters were conducted by CAT, which resulted in 21.98% (20/91) and 4.39% (4/91) of them assigned as Proteobacteria and Thaumarchaeota, respectively (Fig. 5A and Table S7). As a dominant primary producer in deep-sea water layers, Thaumarchaeota have several distinct lineages of ammonia-oxidizing archaea in deep ocean ^54^. We demonstrate the distribution of their AMP clusters by calculating CPM in the five depth zones. The AMP RfSP clusters k11_399729_1 (cluster_479) identified from the SCS metagenomes was almost restricted in shallow epipelagic, mesopelagic and bathypelagic zones (Fig. 5B). However, the two clusters M11_37_77 (cluster 31381) and M11_70_49 (cluster 55418) were derived from MT metagenomes and were abundantly present only in abyssopelagic and hadopelagic zones, while were absence in epipelagic or mesopelagic zone (Fig. 5B). Most of them were unknown in function with respects to CDD and EggNOG annotations, except for a domain similar to PyrH (UMP kinase).

**Figure 5.**
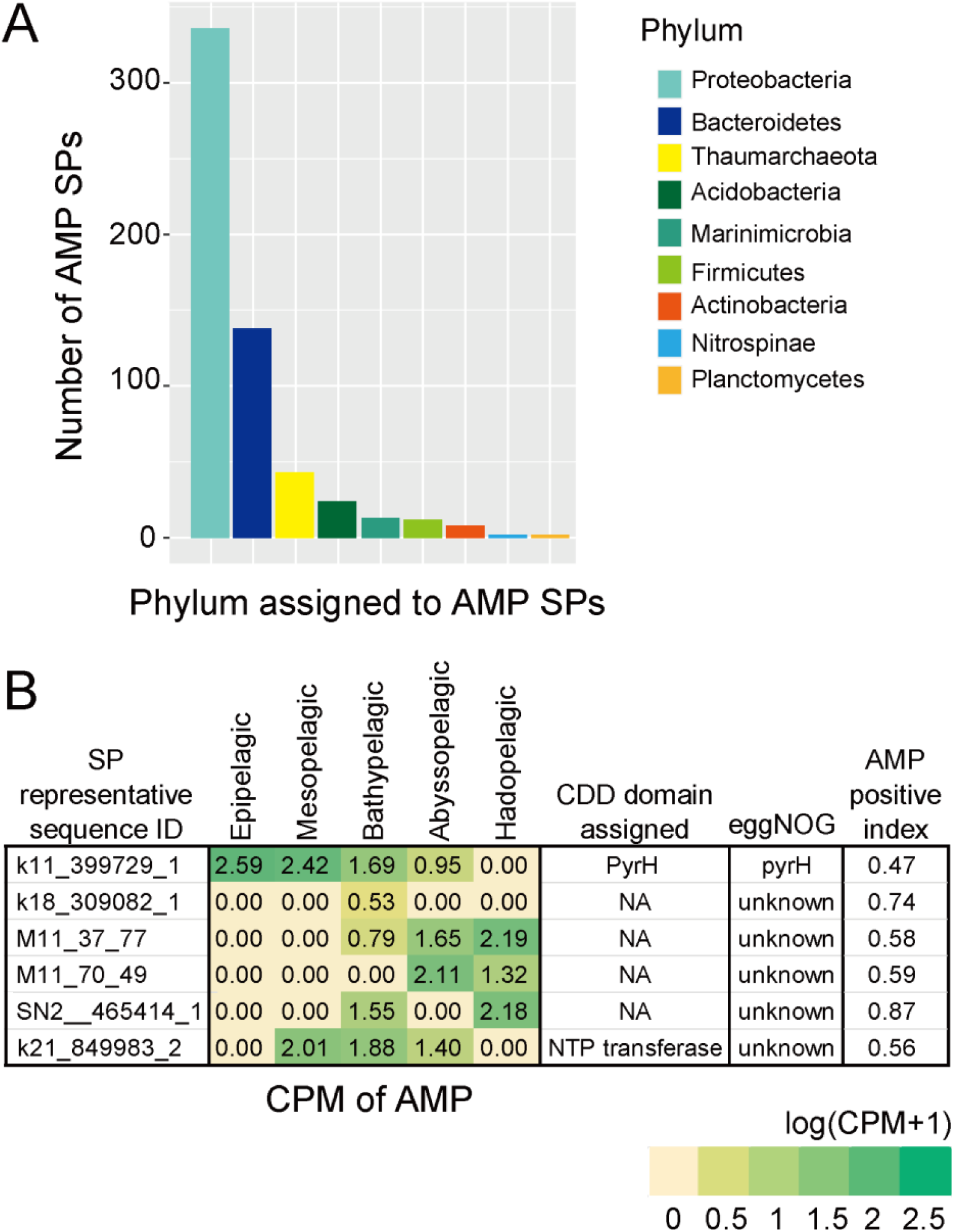
Predicted antimicrobial RfSPs. (A) Taxonomic assignment of the predicted AMPs at phylum level; (B) Average CPM in each depth zone, functional description (CDD and EggNOG), and positive index predicted by AMPfun (Chung, C. R., *et al*, 2020) for the five AMPs derived from Thaumarchaeota. CPM: copy number of sORF per million reads.

### Detection of small proteins in metatranscriptomes and metaproteomes

To examine transcriptional level of the SP clusters, we mapped the representative sORF sequences of 4,307 RfSP and 75,581 prevalent SP clusters to 13.16 Gbp clean reads of 12 metatranscriptomes of water samples that were collected with a multiple *in situ* nucleic acids collection (MISNAC) equipment from a ∼1,000 m depth site in SCS ^55^. The result showed that 3.50% (2,605/77,581) of representative sequences of the prevalent SP clusters could be detected in the metatranscriptomes, while 8.20% (353/4,307) of the RfSP clusters were transcribed. 97.17% (343/353) of the transcribed RfSP clusters overlapped with the transcribed prevalent (RNAcode filtered) SP clusters.

The number of transcribed RfSP clusters detected in the mesopelagic (n=298) and bathypelagic zones (n=313) was higher than other zones (Fig. 6A). This is likely due to the mesopelagic source of the metatranscriptomes. About 30.32% (104/343) of the transcribed RfSP clusters lack an RBS motif, indicating transcription without a ribosome binding site or in a manner lacking 5’-UTR as in eukaryotes ^32^. 5’GGA/GAG/AGG’3 was the most preferred RBS spacer used by the sORF members (Fig. 6B and Table S8). 48.33% (1172/2,425) transcribed RfSPs were classified as Proteobacteria and 20.37% (494/2,425) of them were classified as Thaumarchaeota (Fig. 6C and Table S8). In the transcribed RfSPs, the percentage of transmembrane or secreted proteins in mesopelagic (on average 30.47%) and bathypelagic (on average 33.11%) zones was higher than those in other zones (Table S8), which indicates that these SPs might play a vital role in communication among microbes, especially in mesopelagic and bathypelagic zones (Fig. 6D and Table S8). Among the transcribed RfSP clusters, 231 had an EggNOG annotation description and some of them might contribute to cross-membrane transport (n=23), protein folding (shock protein) (n=9), cell division (n=3) and DNA stability (n=15) for deep-sea adaptation (Table S8). Some known heat-shock proteins as molecular chaperones are constituted by few amino acids ^56^. These may give a clue that chaperone SPs were associated to ‘cold-shock’ response in deep-sea prokaryotes as well. Two SP clusters were described as components of proline/glycine betaine transporter (Table S8). RfSP clusters 114326 (glycine betaine L-proline transport ATP binding subunit) and 30184 (Substrate binding domain of ABC-type glycine betaine transport system) may enhance the up-taking glycine betaine transporter as the mimic function of AcrZ in AcrAB-TolC efflux pump ^57^. These two SP clusters might help prokaryotes to transport glycine betaine to adapt to high hydrostatic pressure in deep sea ^58^. RfSP clusters 4092 and 1795 assigned to Thaumarchaeota were annotated as ammonium transporter (Table S8), which is a membrane-integrated protein ubiquitous in microorganisms for uptake of ammonia for subsequent amino acids biosynthesis and ammonia oxidation ^59^. Furthermore, the transcribed RfSP cluster 16132 was predicted to be an ATP-dependent serine protease Lon that mediates the selective degradation of mutant, aberrant and certain short-lived regulatory proteins ^60^, capable of degrading polypeptides progressively to short peptide fragments ^61^ that are required to rapidly maintain cellular homeostasis and resist stress-induced DNA damage ^62^. This suggested that SP cluster 16132 might help prokaryotes adapt to rapidly changing deep-sea environments such as hydrostatic pressure. The RfSP clusters 55310, 40899 and 52138 were involved in cell division (Table S8). The SP cluster 3931 (HupA) (Table S8), a histone-like DNA-binding protein, was detected in our metatranscriptomes. The HupA which is able to encapsulate DNA inside to achieve DNA sequence stabilization ^63^, thereby preventing DNA from denaturation under high hydrostatic pressure^64^. This SP cluster was abundant in the metagenomes for the water samples collected in bathypelagic zone (1,000-4,000 m depth), suggesting that it may help prokaryotes to maintain DNA stability and resist extreme environments in the deep ocean.

**Figure 6.**
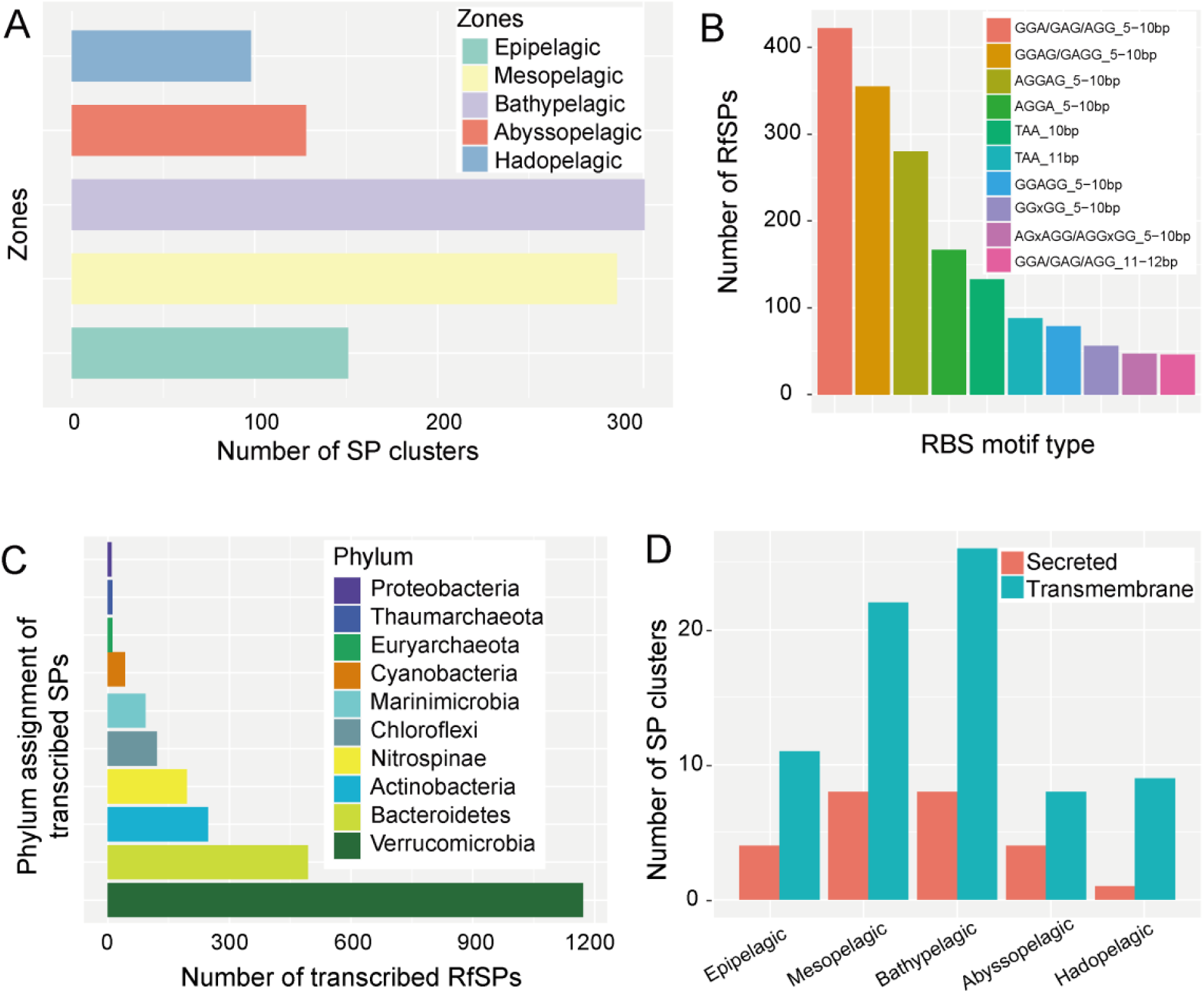
Features of transcribed RfSPs[1]. (A) Number of RfSP clusters present in each depth zone; (B) Number of 5∼10-bp RBS spacers used by transcribed RfSP clusters; (C) Taxonomic classification of transcribed RfSPs at phylum level; (D) Number of transmembrane/secreted RfSPs in the transcriptomes (details in Table S9).

To confirm that the SPs were translated, we searched the SP clusters in 14 metaproteomes of deep-sea samples from SCS and MT ^65^. A total of 38 SP sequences among 75,581 prevalent SP clusters have homologs with >60% amino acid identity and >60% sequence coverage in the metaproteomes. The results showed that four SPs were associated with an assignment of conserved domain (Table S9). Among the translated SPs, SP cluster 40295 containing a domain assigned as acetolactate synthase, which is the first enzyme playing divergent roles in branched-chain amino acid synthesis ^66^. The translated SP cluster 74952 might be superoxide dismutase. One of the two translated SPs that could be annotated by the EggNOG database was probably a cold shock protein. Hence, both metatranscriptomes and metaproteomes showed that SP might play an important role in hyperoxide detoxification and protein-folding correction. With more deep-sea samples for proteomics work, a high ratio of the SPs of this study will be shown in metaproteomes.

## Conclusions

In this study, we report the SPs from the full-ocean-depth marine water samples and partially confirm their transcriptional and translational activities. About 70% of 75,581 prevalent SP clusters lack a functional description, which indicates that marine water column is a vast reservoir of novel SPs for screening new efficient gene editing tools. The known functions of the SP clusters suggest that they play a particularly vital role in altering genome structure, resisting hydrostatic pressure, defending against viruses, and rapidly responding to changes in the environment for the dominant deep-sea species. Our study shows a high proportion of AMP SPs from Archaea, indicating a goldmine of antibiotics in the archaea from the dark ocean. Future work will validate the conclusions of this study regarding the prevalence and distribution in water layer and species with meta-omics data from global oceans.

## Materials and methods

### Sample collection, DNA extraction and sequencing

Sixty-nine marine water samples were collected from MT and SCS at depths in a range of ∼100 m to 10,909 m during several cruises including *R/V* DY37II, *R/V* TS03, *R/V* Tan Kah Kee and *R/V* Haigong623 (Table S1). About 20 L water samples collected by Niskin bottles were filtered through a 0.22-μm polycarbonate membrane and stored in −80 °C until use. The samples from the hadal layer (>6,000 m) were obtained by an *in situ* sampler ISMIFF with capacity of filtration and fixation ^42^. The polycarbonate membranes were cut into small pieces for DNA extraction using MO BIO Power Soil DNA Isolation Kit (MoBio, Carlsbad, CA, USA) according to the manufacturer’s instruction. The DNA/RNA coextraction samples collected by MISNAC for generation of the SCS metagenomes were described in details in our previous study ^55^. The quality and quantity of the DNA extraction were examined by 1% agarose gel electrophoresis and Qubit 2.10 fluorometer (Invitrogen, Life technologies, USA). The good-quality DNA was first sheared randomly to fragments of ∼500 bp or 350 bp by Covaris Focused-ultrasonicator M220 (Covaris, LLC., Woburn, Massachusetts, USA) and used for metagenomic library preparation with TruSeq® Nano DNA LT Sample Prep Kit (Illumina, San Diego, CA, USA). The high-throughput sequencing was performed on an Illumina Miseq platform (2 × 300 bp) or a Novaseq 6000 platform (2 × 150 bp). Two additional metagenomes were obtained for the Mariana surface water (MT0m) and 203-m depth water (MT203m) from the NCBI database (accession number PRJDB5777).

### Raw data processing, de novo assembly and prokaryotic contig selection

The metagenomic raw data were filtered by Fastp (v.0.20.0)^67^. The clean reads after removal of replicate sequencing reads by Fastuniq ^68^ were assembled with SPAdes (v.3.13) ^69^(kmer set of 21, 33, 55, 77, 99 and 127) or MEGAHIT^70^. Eukaryote contigs were removed using EukRep with strict model (-m strict)^71^.

### Identification and filtration of SP clusters

The ORFs in all the clean prokaryotic contigs were predicted by Prodigal (v.2.6.3) ^72^ with -meta model. The shadow ORFs (an ORF that substantially overlaps with an aORF and forms a potential overlapping gene pair) were identified and removed with an R script provided by a previous report ^73^. Spurious ORFs were identified by hmmsearch (v.3.2.1) ^74^ against AntiFam database ^75^ and then removed with an e-value cutoff of 1e-05. The complete ORFs (including start and stop codons) in size of ≤150 bp were retained. The deduced amino acid sequences of the ORFs were clustered by CD-HIT (v4.6.8) (-c 0.5; -s 0.95; -aL 0.95; -g 1; -n 2; -p 1; -d 200; -M 50000; -l 5). The clusters were further checked according to the following steps: 1) building a Blastn database of the representative sORF sequences of all the SP clusters; 2) the clean reads of the metagenomes were queried to the sORF database; 3) the mapped reads with 90% coverage and 95% identity were recruited for CPM calculation; 4) the prevalent clusters with a CPM value ≥5 for any of the metagenomes were retained; 5) All the SPs of a prevalent cluster that no less than three sORF sequences were treated by MAFFT (v. 7.471) ^76^ for further filtered by RNAcode (v.0.3)^30^ with *p* ≤ 0.05; 6) the RfSP clusters in which all the sORFs were present at the last position of the respective contigs were removed to filter premature ORFs caused by sequencing errors.

### Taxonomic assignment of SP clusters

The CAT database was downloaded from https://tbb.bio.uu.nl/bastiaan/CAT_prepare. The prokaryotic contigs containing any sORF in the RfSP clusters were identified and classified by CAT ^39^ with contigs model.

### Annotation of small proteins

The representative sequences of RfSP clusters were annotated by eggNOG-mapper (v.2.0.0) with eggNOG database (v.5.0) ^77^(--seed_ortholog_evalue 0.00001; --query-cover 90; --seed_ortholog_score 60) and searched against the NCBI_Nr database with DIAMOND Blastp (v.2.0.11.149) ^78^(--query-cover 90; --sensitive; --evalue 1e-5) with identity threshold of 95%. The reviewed (true) protein sequences with length less than 200aa were downloaded from UniPort (January, 2024) for a self-built database, and representative sequences of RfSP clusters were searched to it by BlastP (evalue, 1e-5; length of hit spans 90%∼110% of representative SP).

All amino acid sequences of 75,581 prevalent clusters after CPM filtration were annotated by RPS-BLAST^79^ (e-value 1e-02; coverage 80%) against CDD (September. 2022; v.3.20)^80^. Position Specific Scoring Matrices (PSSM) IDs of ribosomal proteins were collected from CDD by key word “ribosomal protein” and checked manually (Table S4). The nonredundant PSSM IDs were searched in the CDD annotation result of all SP sequences in the prevalent clusters.

All prokaryotic genomes were downloaded from the GenBank of NCBI database (https://ftp.ncbi.nlm.nih.gov/refseq/release/bacteria/) (November, 2023; released version 221) and all ORFs were predicted by Prodigal (v.2.6.3) ^72^. Then, ORFs coding for no more than 50 aa were picked to build a self-curated database. The representative sequences of RfSP clusters were searched against the database by BlastP (v2.2.29) (-evalue 1e-5, -word_size 2, -max_hsps 500). The representative sequences of the RfSP clusters were searched against the database including proteins with *≤*50 aa from prokaryotic protein files in NCBI with the same parameters. Finally, all the search results were filtered with thresholds as following: length of the hit was 90% ∼ 110% of the length of the representative SP.

### Identification of SP clusters potentially involved in cell communication

Signal peptides of all amino acid sequences of the RfSP clusters were predicted by local signalp-5.0 (-f summary -t gram--c 0; -f summary -t gram+ -s no TM -u 0.44 -c 0)^81,82^. An SP cluster was recorded if more than 80% of the amino acid sequences contained a signal peptide (secreted). Transmembrane structure of SPs in each cluster was predicted using tmhmm (v.2.0)^83^ with the same criteria as those for secreted peptides. Phobius (v.1.01)^84^ was also used for prediction of signal peptide and transmembrane topology of SPs in each cluster.

### Prediction of antimicrobial peptides among SP clusters

AMPs in the RfSP clusters were identified by searching their representative amino acid sequences in Anti-Microbial Peptides (CAMP_R3_) ^85^(http://www.camp.bicnirrh.res.in/prediction.php) and Database of Antimicrobial Activity and Structure of Peptides (DBAASP) (v.3.0) ^86^(https://dbaasp.org/prediction/general). During the AMP prediction by CAMP_R3_, all algorithms were used including support vector machines (SVM), random forest (RF), artificial neural network (ANN) and discriminant analysis (DA) for screening positive AMPs. An AMP SP must be positive in both CAMP_R3_ (confirmed by four types of CAMP_R3_ algorithms) and DBAASP. The AMP activities against specific microbial species were predicted in DBAASP.

### Prediction of small proteins related to defense and HGT

The functions of SP clusters were further inferred from the CDD annotation results of 10 up-/down-stream sORFs in the contig where the sORF encoding the representative SP was located. To recruit genes related to defense systems, 427 known genes involved in defense system were collected as demonstrated by a previous study ^51^. In addition, 35 genes of all known CRISPR-Cas systems were also added into the list, including the words of “CRISPR”, “Cas”, “tracrRNA” (transactivating CRISPR RNA) and ancillary genes of CRISPR-Cas system except “TPR” as summarized previously ^50^. All the genes were used to search against CDD annotation results of all the ORFs in the same contig containing a corresponding representative sORF.

### Searching the marine SPs in non-marine metagenomes

The contigs of eighty non-marine metagenomes were downloaded from JGI (Joint Genome Institute) or MGnify (former EBI Metagenomics) ^87^. The representative DNA sequences of the RfSP clusters were searched against all the contigs of 80 non-marine metagenomes by Blastn ^79^ with several parameters (e-value, 1e-05; identity ≥50%; coverage ≥90%). Then ORFs in the filtered contigs were predicted and were used as a database. The representative sequences of the RfSP clusters were queried against it by Blastp ^79^ with e-value cutoff of 1e-05 and length of hit ranging from 90% to 110% of that of RfSP representative.

### Transcriptional activity of SPs in deep-sea metatranscriptomes

The nucleic acids of 12 marine water samples were collected *in situ* with the MISNAC equipment at ∼1,000 m depth of SCS as mentioned above ^55^ (Table S1). The metatranscriptomes were generated as described in our previous study^88^. The rRNA gene reads were removed by SortMeRNA (v4.3.4) from the metatranscriptomes ^89^. The clean reads of metatranscriptomes were mapped to the representative sequences of the SP clusters by Blastx with a coverage threshold of 50%, identity ≥90% and e-value <10^−5^. TPM was calculated as previously described by taking read length and gene mapping rate into consideration ^26,90^.

### Identification of translated SPs in marine metaproteomes

The metaproteomics data of marine water samples collected from the Northwest Pacific Ocean (Fig. 1) were downloaded from ProteomeCentral (ftp://ftp.pride.ebi.ac.uk/pride/data/archive/2021/03/PXD014630)^65^. The deep-sea surface sediments from the Mariana Trench for metaproteomes TS03_B11_10900 and DY37_DIVE120 were collected by *R/V* TS03 cruise in 2017 and *R/V* DY37II cruise in 2016. The marine sediments were treated with TRIzol Reagent (Invitrogen^TM^, Carlsbad, CA, USA) for protein extraction according to the manufacturer’s protocol. The organic phenol-chloroform phase and interphase were subjected to a protein isolation procedure; finally, 100 μL of 10 M urea with 50 mM DTT (Sigma, St. Louis, MO, USA) was used to dissolve the protein pellet, followed by a centrifuge at 16,000× g for 10 min. The supernatant was then collected into a new tube. 40 mM iodoacetamide (Sigma) was used for protein alkylated and 25 mM tetraethylammonium bromide (TEAB; Sigma) for dilution. Trypsin (Promega, Madison, WI, USA) was used to protein digestion with a protease:protein ratio of 1:50 (w/w), followed by incubation for 16 h at 37 °C. After tryptic digestion, LC-MS/MS detection was performed ^91,92^. All the protein sequences of the metaproteomes were queried to representative sequences of 75,581 prevalent SP clusters by Blastp (v.2.13.0) with criteria of coverage percentage ≥60%, identity ≥60%, and e-value <10^−5^.

### Availability of Data

SAR data of two samples (MT_0m, MT_203m) were downloaded from national center for biotechnology information (NCBI) (PRJDB5777). The prokaryotic SPs had been deposited in the National Omics Data Encyclopedia (NODE) with OEP004991 as project ID (https://www.biosino.org/node/project/detail/OEP004991) and OER455278 as run ID (https://www.biosino.org/node/run/detail/OER455278). The contigs of this study for SP identification were deposited in the database archive (https://www.biosino.org/node/run/detail/OER455278#ftp-download). The RfSP clusters are accessible from the database with the link: https://www.biosino.org/node/run/detail/OER455278#ftp-download.

In “contig_sorf.fa” file, “>k10_contig_227_fa” means the 227th contig of sample named k10.

In “sp4307_orf.faa” file, “>M5_contig_19873_fa_3_faa” means that the sORF is the third ORF of the 19873rd contig of sample named M5.

## Funding

This study was supported by the National Natural Science Foundation of China (42376149), and the Shenzhen Key Laboratory of Advanced Technology for Marine Ecology (ZDSYS20230626091459009).

## Author contributions

Q.M.L. conceived the work, performed the analysis and wrote the manuscript; Q.M.L., L.S.H. and Y.W. designed the study; L.S.H. and Y.W. critically revised this manuscript; All the authors read and approved the final manuscript.

## Acknowledgements

We also thank the supercomputer center of Sanya University. This research was supported by the National Natural Science Foundation of China (No. 42376149).

## Conflict of interests

The authors declare that there is no conflict of interests.

## Notes

### Competing Interest Statement

The authors have declared no competing interest.

