## Supplementary material for "Small proteins from prokaryotes in marine water column at full ocean depth": Fig.S1

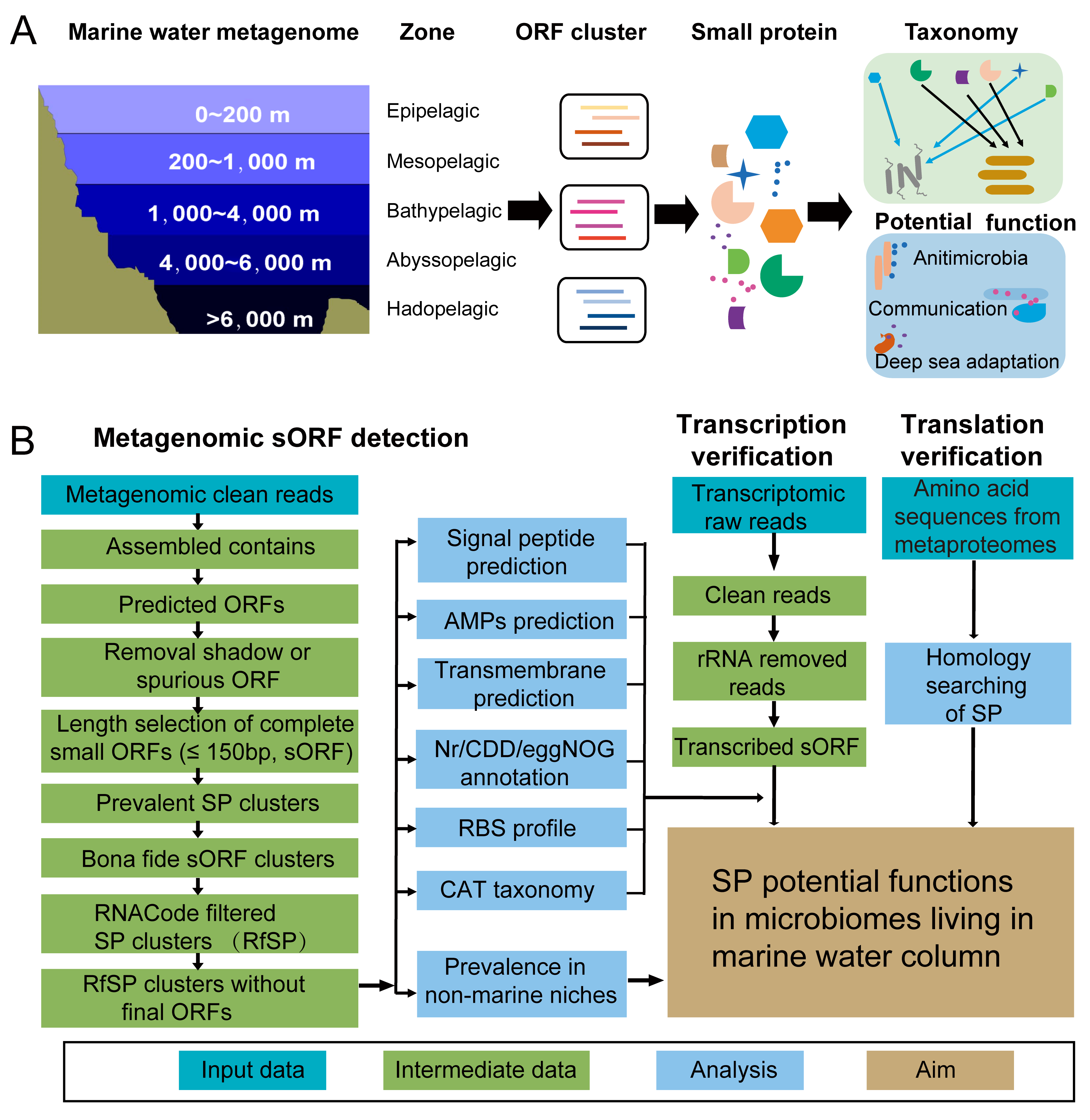

**Figure S1. Workflow illustrating identification and functional annotation of prokaryotic small protein (SP) in marine waters.**

Full-ocean-depth water samples were obtained from the South China Sea and the Mariana Trench. A total of 71 metagenomes were individually assembled for prediction of open reading frames (ORFs) from the contigs. Complete short ORFs (sORFs) encoding no more than 50 amino acids were further recruited and a non-redundant sORF set was built by clustering the sORFs with CD-HIT. The sORF clusters were filtered by ≥5 copies per million reads (CPM) in all samples to identify *bona fide* sORFs. The potentially transcribed sORF clusters were predicted by RNAcode (Washietl, S. *et al*., 2011). The annotation of small proteins (SPs) was performed by running RPS-BLASTP, diamond blastp, and emapper queries against CDD, NCBI-nr, and eggNOG databases, respectively. The taxonomy of the contigs encoding SPs was assigned by CAT (von Meijenfeldt, F A Bastiaan *et al.* 2019). The prediction of secreted, transmembrane, and antimicrobial SPs was performed by signalP and TMHMMhmm. Transcriptional activity of the SPs was confirmed by retrieving *in-situ* metatranscriptomic reads that mapped to the corresponding sORFs. These sORF clusters were queried to the protein sequences of fourteen metaproteomes of marine waters and surface sediments.

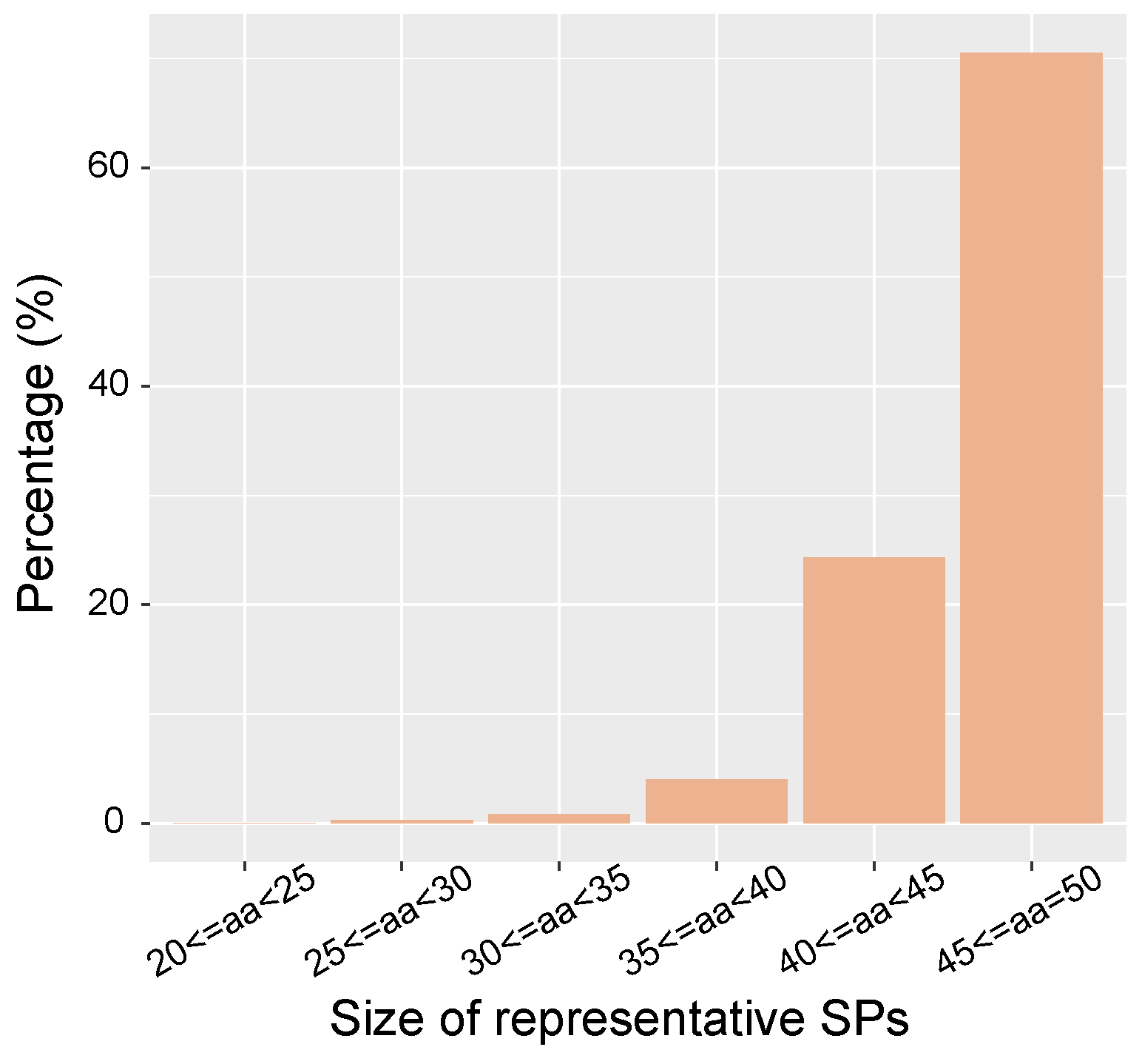

**Figure S2. Length distribution of representatives of 4,307 RNAcode-filtered SP clusters from full-ocean-depth marine prokaryotes.**

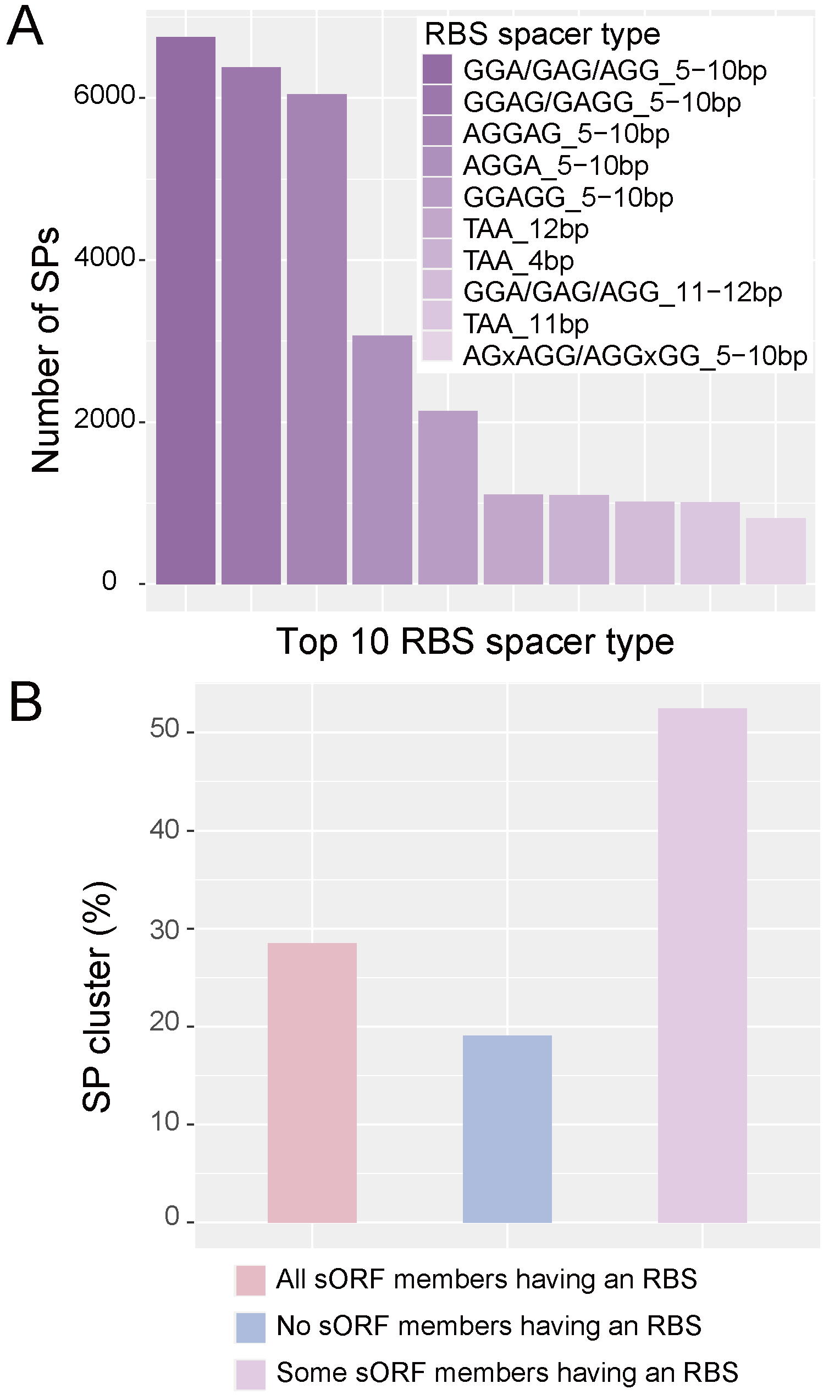

**Figure S3. Characteristics of ribosomal binding sequence (RBS) motif of 4,307 RNAcode-filtered SP clusters from full-ocean-depth prokaryotes.**

(A) The top 10 most frequent spacers of ribosomal binding site (RBS) motif in the sORF (B) Percentage of SP clusters with/without/partially with a ribosomal binding site (RBS) in the sORF members.

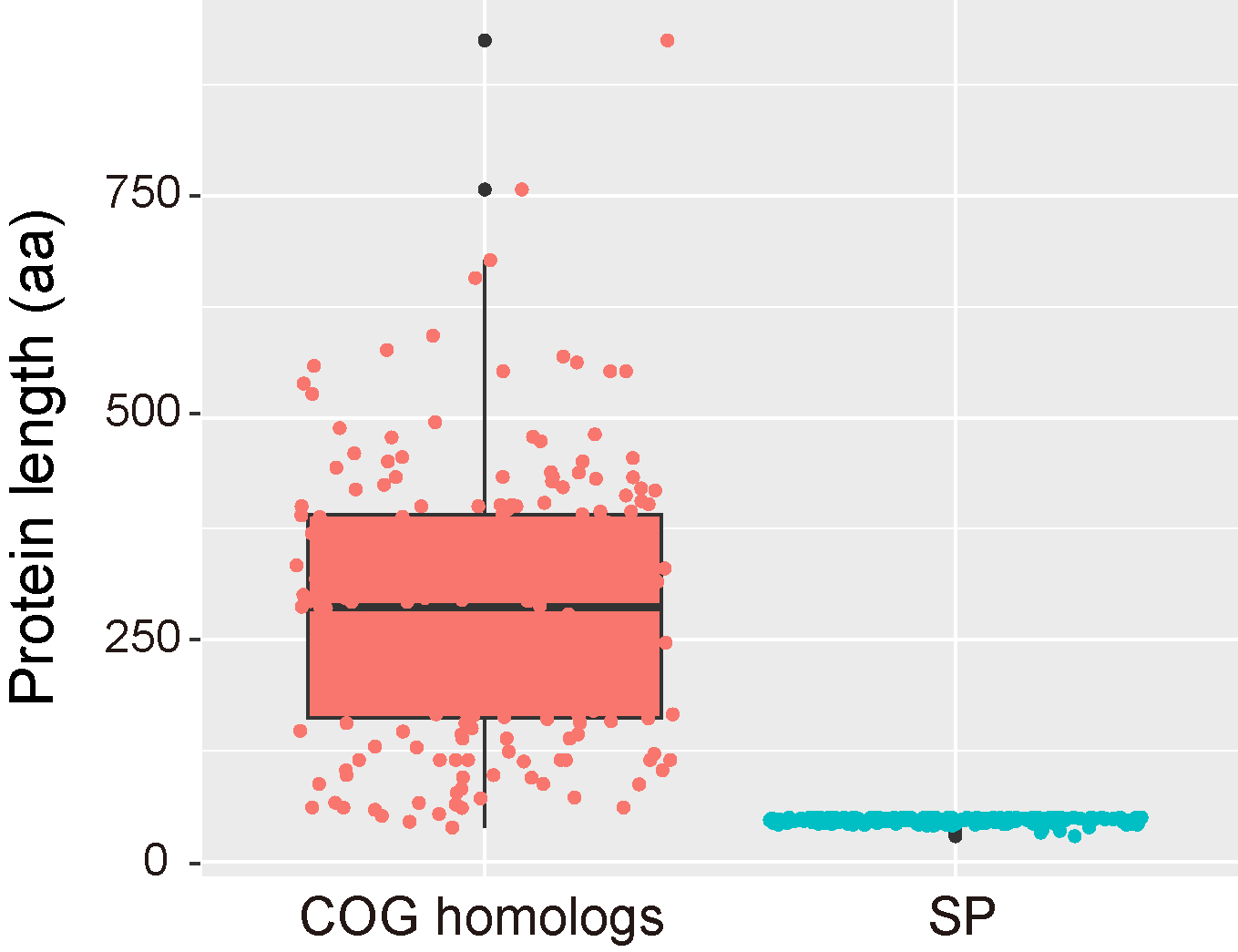

**Figure S4. Length distribution of SPs and their homologies in COG database.**

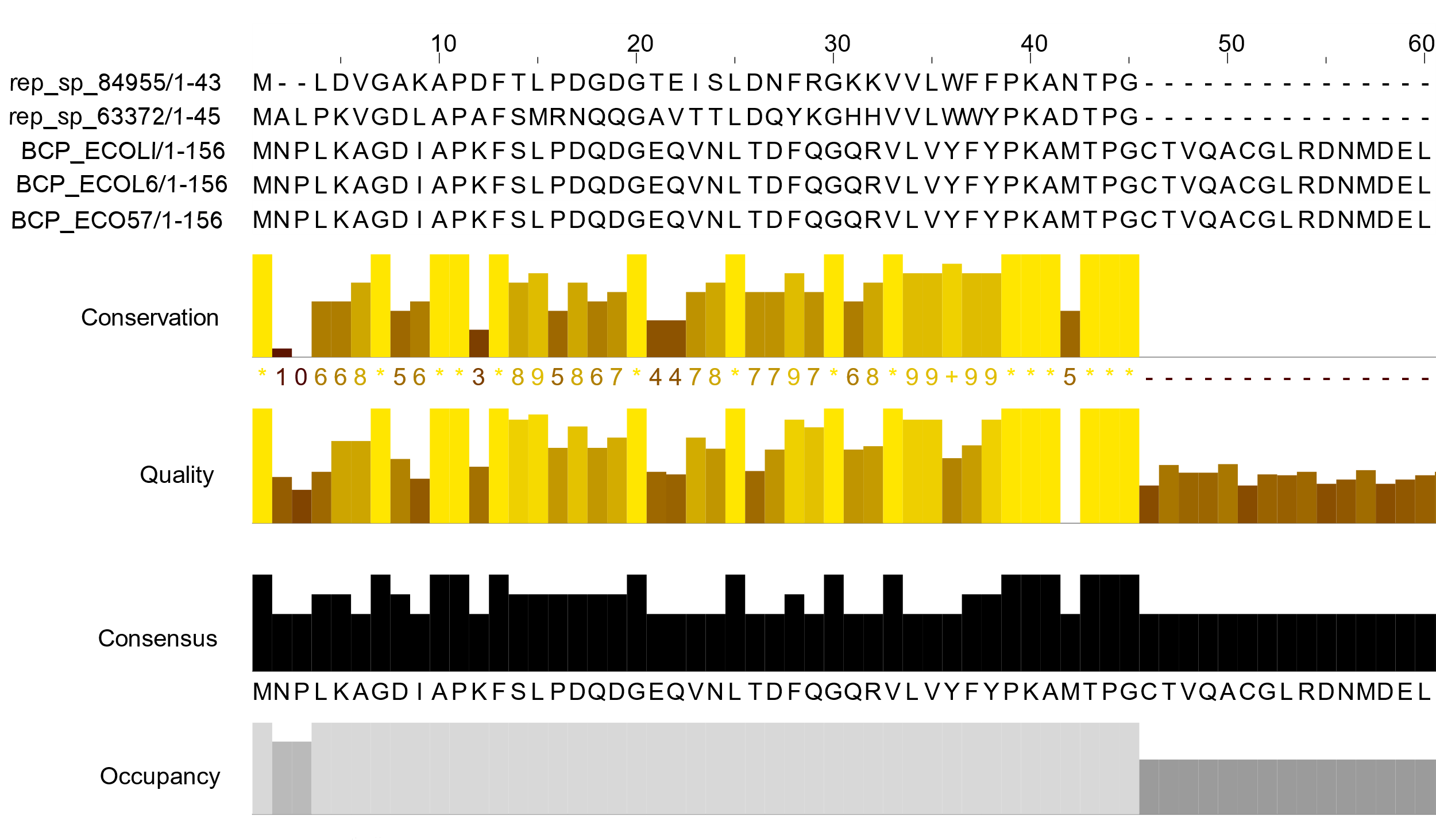

**Figure S5. The alignment of representative BCP SPs and long homologs of *Escherichia coli*.**

The amino acids sequence alignment and conservative assessment of BCPs were performed for representative SP sequences of clusters 84955 and 63372. The first 60 amino acids were shown for the reference BCPs of *E. coli* from the Uniport database.

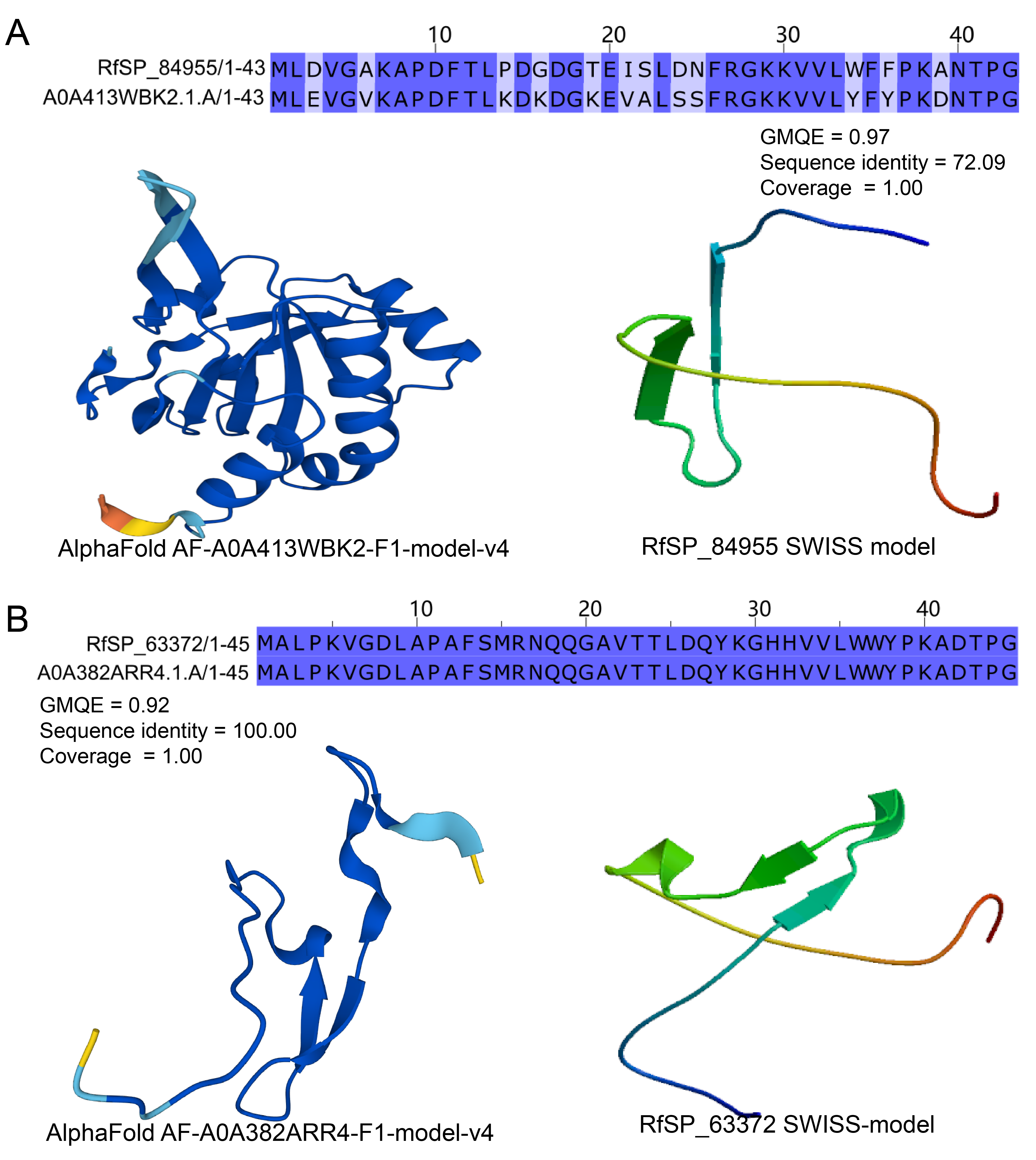

**Figure S6. Structural similarity of representative SPs assigned as BCP proteins.**

The structural prediction was performed using representative SP sequence of RfSP clusters 84955 and 63372.
